# Strongly correlated spatiotemporal encoding and simple decoding in the prefrontal cortex

**DOI:** 10.1101/693192

**Authors:** ED Karpas, O Maoz, R Kiani, E Schneidman

## Abstract

We studied the fine temporal structure of spiking patterns of groups of up to 100 simultaneously recorded units in the prefrontal cortex of monkeys performing a visual discrimination task. We characterized the vocabulary of population activity patterns using 10 ms time bins and found that different sets of population activity patterns (codebooks) are used in different task epochs and that spiking correlations between units play a large role in defining those codebooks. Models that ignore those correlations fail to capture the population codebooks in all task epochs. Further, we show that temporal sequences of population activity patterns have strong history-dependence and are governed by different transition probabilities between patterns and different correlation time scales, in the different task epochs, suggesting different computational dynamics governing each epoch. Together, the large impact of spatial and temporal correlations on the dynamics of the population code makes the observed sequences of activity patterns many orders of magnitude more likely to appear than predicted by models that ignore these correlations and rely only on the population rates. Surprisingly, however, models that ignore these correlations perform quite well for decoding behavior from population responses. The difference of encoding and decoding complexity of the neural codebook suggests that one of the goals of the complex encoding scheme in the prefrontal cortex is to accommodate simple decoders that do not have to learn correlations.

## INTRODUCTION

The analysis of neural codes has often been described in terms of characterizing two “dictionaries:” An encoding dictionary that maps stimuli into neural responses and a decoding one, which maps neural activity into estimated stimuli or behavioral outcomes [1, 2]. While encoding and decoding are strongly coupled, their respective dictionaries may have very different structures and properties. In engineering, encoding is typically designed to maximize information rate through overcoming noise, whereas decoding aims at reconstructing the original signal, minimizing a well defined error function [3]. The brain operates under very different requirements and constraints. Neural encoding needs to convey the relevant information to downstream circuits, in an efficient way under constraints of time, energy, noise, and number of neurons [2, 4, 5]. The nature of the code also depends on the architecture and state of the encoding network of neurons, which may create complex and even variable codes in different encounters with the same stimulus or execution of similar actions, especially for neural processes that evolve over time [6]. Neural decoding needs to give a reduced representation of the stimulus, often in terms of a decision or action for the organism to take, whereby minimizing an error function that is typically unknown in full to us. Also, what information is relevant may vary from one moment to the next, depending on the needs and goals of the organism. Thus, decoding in the brain is typically far from a direct inverse of encoding and would typically retrieve only part of the information encoded in upstream networks; in engineering terms this would mean a very “lossy” decoding [3]. So, despite their inherent coupling, neural encoding and decoding could follow different operational principles.

We study here the nature of the population code of the prefrontal cortex (PFC) in monkeys during a perceptual decision making task. The PFC is an especially interesting brain region to study population encoding and decoding due to its role in many high-level cognitive functions [7–9], and the prevalent recurrent connectivity and large number of synapses between its neurons [10, 11], which provide a structural basis for rich dynamics to emerge. Decision making is typically a temporally extended process, often described in terms of different functional stages: gathering information, transforming information to a choice, and acting upon the choice [12]. Accordingly, the relation between neural activity and decision making is commonly studied using tools from decision theory, e.g., by projecting neural activity on a decision variable [13–16]. In particular, for perceptual decisions based on accumulation of sensory evidence, it has been shown that dynamics of population firing rates reflect the accumulation process that leads to the decision [13, 16, 17].

Characterization of encoding in large neural populations hinges on mapping the spatial and temporal structures of spiking patterns, and how they carry information [18–24]. Past analyses have largely relied on the firing rates of individual units and their dynamics. Yet, in many neural circuits, the joint activity patterns of large populations of neurons have been shown to be strongly correlated on short time scales [25–28], both in terms of signal correlations [29–33] and noise correlations [34–36]. Thus, the study of population codes requires a detailed mapping of the joint spiking patterns of large populations of neurons and their evolution through time, without making a priori assumptions, to uncover the underlying design principles in terms of spiking patterns, correlations, time scales, and their functional implications. We therefore use statistical modeling of population spiking activity at fine temporal resolutions to characterize the spatial and temporal structure of population dynamics and their functional timescales in the prefrontal cortex of monkeys.

Using highly accurate statistical models of the spiking patterns of 100 units, we show that strong spatial and temporal correlations at the population level shape the encoding dictionary. We further characterize history-dependence and the dynamics of population spike patterns, demonstrating that the spike patterns and their sequences strongly depend on the epoch of the task and computations performed by the network. The timescale of temporal dependence in the code is only tens of milliseconds, suggesting that conventional rate-based analyses that rely on longer analysis windows would fail to accurately capture the neural code. Finally, we show that despite the strong spatio-temporal correlations in encoding, and maybe because of them, the resulting code is one that is readable by a simple decoding scheme that does not take these correlations explicitly into account. These properties make the neural code suitable for versatile and complex computations while also making it easy for downstream areas to interpret prefrontal responses.

## RESULTS

We recorded the activity of groups of over 100 units from the prefrontal cortex of two macaque monkeys performing a perceptual decision making task. Electrophysiological recordings were done using Utah arrays, over 15 sessions, each consisting of more than a thousand trials (see Methods). Monkeys were trained to perform a direction discrimination task based on the movement of random dots [13, 37]. On each trial, the monkey fixated on a central fixation point, and then presented with two targets representing the two possible motion directions. After a short interval, a patch of randomly moving dots was shown, followed by a delay period of variable length. After the delay, the monkey made a saccade to one of the two targets, based on the net motion direction (Fig. 1A), and then maintained fixation on the chosen target until feedback. Correct choices were rewarded with juice, and errors were signaled with a distinct tone. As previously reported, monkeys’ performance was strongly correlated with task difficulty, ranging from chance for 0% motion coherence to perfect accuracy for the highest motion strength [17, 38, 39]. Further, single cell responses varied both with motion strength and choice [16, 17, 38].

**Figure 1:**
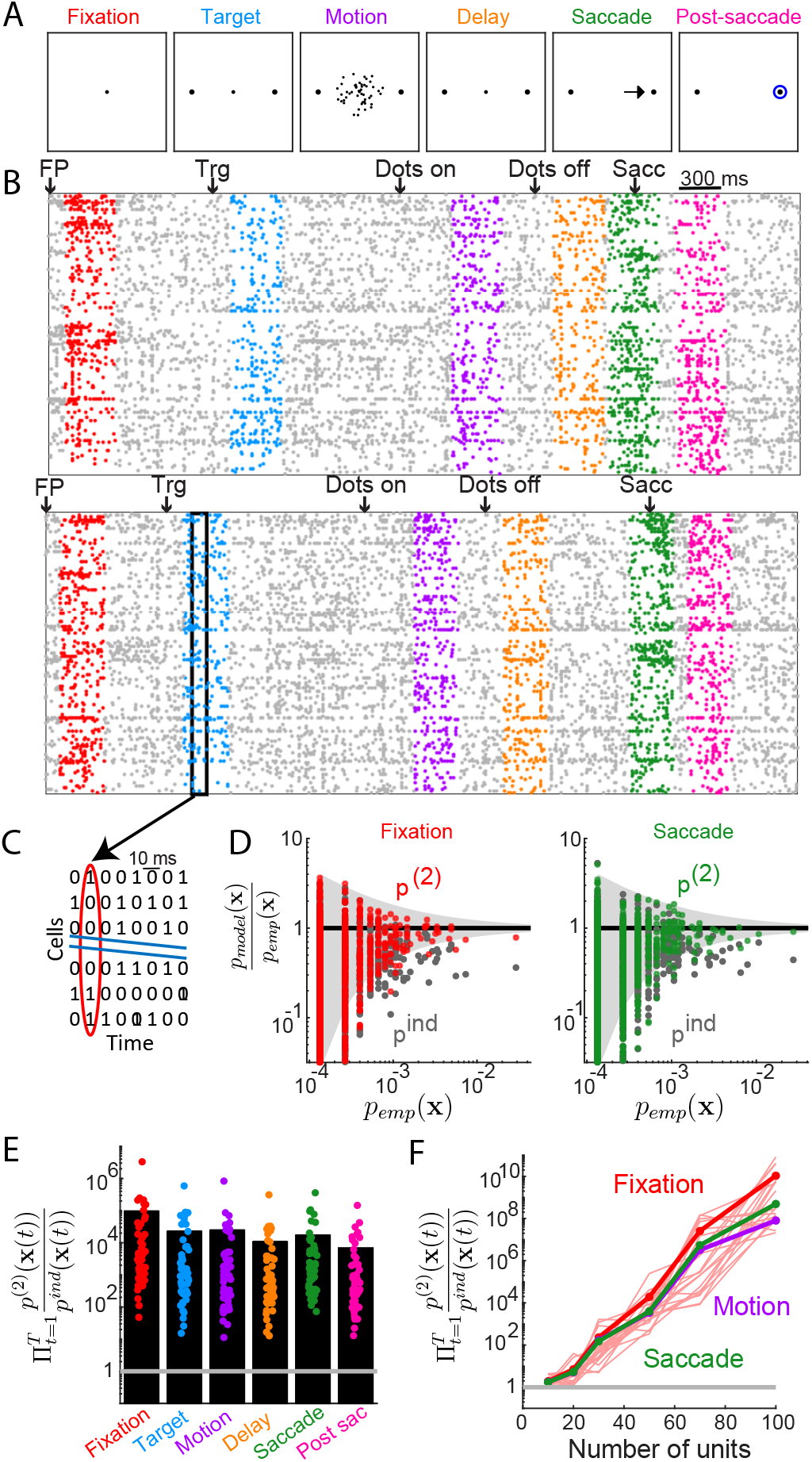
Behavioral task and correlated activity of pre-arcuate gyrus neural populations in different task epochs. **A.** Task design. Sequence of epochs in each trial: Fixation, Target presentation, Dots motion, Delay period, Saccade to chosen target, and Post-saccade fixation (see main text and Methods). To facilitate distinction of epochs throughout the paper, a unique color is designated to each epoch. **B.** Examples of population spiking raster plots from two trials where the monkey chose the left (top) or the right target (bottom). Analyses focused on 300 ms windows in each of the six task epochs, as marked by the corresponding epoch color. Particular events during the trial - appearance of fixation point (FP), and of target (Trg), etc. are marked by arrows above spike rasters. **C.** Example of spike patterns after binary discretization into 10 ms bins. **D.** Examples of predictions of the independent and pairwise maximum entropy models for observed population spiking patterns of a group of 50 units in two epochs: fixation (left) and saccade (right). The gray funnels show the 99 % confidence interval due to the finite amount of data. **E.** Likelihood ratio between the pairwise and independent models in all epochs. Dots mark individual groups of 50 units (randomly chosen), and black bars denote averages over 50 random groups of 50 units. Note that the y axis is a logarithmic scale. **F.** Likelihood ratio between the pairwise and independent models is shown for different population sizes, for 3 representative epochs. Averages over 20 random groups are shown in bold line; values for individual groups are shown for the fixation epoch by thin red lines.

In contrast to previous studies which focused mainly on mean responses of single units or populations of units over extended periods, we set out to characterize spatial and temporal patterns in the spiking activity of the recorded population (Fig. 1B). We discretized the recorded neural activity into Δ*t* = 10 *m*s bins, such that in each time bin, the activity of the population was described by a binary vector x = [*x*_1_, *x*_2_…, *x_N_*], where *x_i_* = 1 designates that the ith neuron spiked in that bin and *x_i_* = 0 means the neuron was silent (Fig. 1C). We analyzed the nature of activity patterns of the population in six non-overlapping task epochs: fixation, target, dots motion, delay, saccade, and post-saccade fixation (Fig. 1A-B). The analyses focused on 300 ms periods in each epoch and, therefore, were done across 30 non-overlapping 10 ms bins.

### Strong correlations among units dominate population vocabulary during all epochs of the task

We start by analyzing the joint activity patterns of the population in single time bins. In all epochs, many of the population activity patterns appeared much more often than expected from the firing rates of the units. Figure 1D shows two examples from different epochs, where models that ignored correlations between units and relied only on the units’ firing rates, failed to predict the prevalence of specific spike patterns. These *independent models*, fitted for each task epoch, are given by 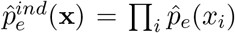, where 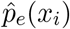 is the probability of cell *i* to spike in each of the time bins in epoch *e*, ignoring the correlations between units. The failure of independent models suggests that the empirical population spiking patterns (vocabulary) demonstrates very strong network correlations, in all epochs, consistent with previous studies of population coding in different neural systems [25, 26, 29, 30, 32, 33].

Because direct sampling of the distribution of population activity patterns for more than a handful of units is impossible — due to the exponential growth of the potential activity patterns with population size — we have to rely on statistical models to quantitatively describe the population code and study it [25, 26, 29, 31]. Since independent models were insufficient, we followed [25, 26, 31], and used the maximum entropy framework to model population activity, taking correlations into account. We recall that for a given set of observable functions of the data, the most parsimonious model among all models consistent with those observables, is the one with the highest entropy [40]. This maximum entropy (ME) model does not make any additional (arbitrary) assumptions, and because entropy is convex, it can be found numerically using gradient descent based methods. We compared the performance of several classes of ME models for the different epochs of our task: the pairwise based models that rely on firing rates of the units and all pairwise correlations between them (*p*^(2)^, which are equivalent to Ising models in physics), the k-Ising models, which extend the pairwise models by adding k-th order synchrony values of the population [31], and random projections (RP) models, which rely on random sets of nonlinear projections of the population activity [32]. Thus, for example, the pairwise models for each of the sessions and individual epoch, *e*, were given by

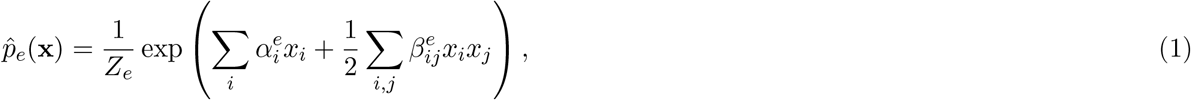

where 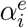 and 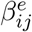 were fit to satisfy the empirical average firing rates, 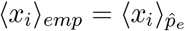 and the correlations between all pairs of units, 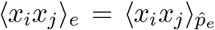. In these equations, 〈〉 denote averages over the empirical data or model, and *Z_e_* is a normalization term or partition function.

We found that the pairwise models were typically orders of magnitude more accurate than the independent models that were based only on firing rates: Fig. 1E shows the comparison of models in one of the sessions, on held out test data for each of the epochs, over many groups of 50 units. On average, the set of observed activity patterns in an epoch were 10^4^ to 10^5^ more likely under the pairwise models than in the independent ones. Similar differences were observed in both animals across different sessions, despite variability for individual sessions and groups of neurons selected for analysis in each session (The median over average groups in sessions was ~ 6 · 10^3^; see see Fig. S1C). The pairwise models, k-Ising models, and RP models were relatively similar in their performance across epochs (see Fig. 2), and so we focus henceforth on pairwise models because of their superior interpretability. Importantly, the collective effect of the correlations among units became larger with population size (Fig. 1F), so that at the level of 100 units, the observed epochs were ~ 10^10^ more likely to appear than expected by independent models. The failure of the independent models and the accuracy of the pairwise models were similar for all epochs in all experimental sessions and animals (Fig. 1C), and the results were robust for time bins of different size (Fig. 7).

**Figure 2:**
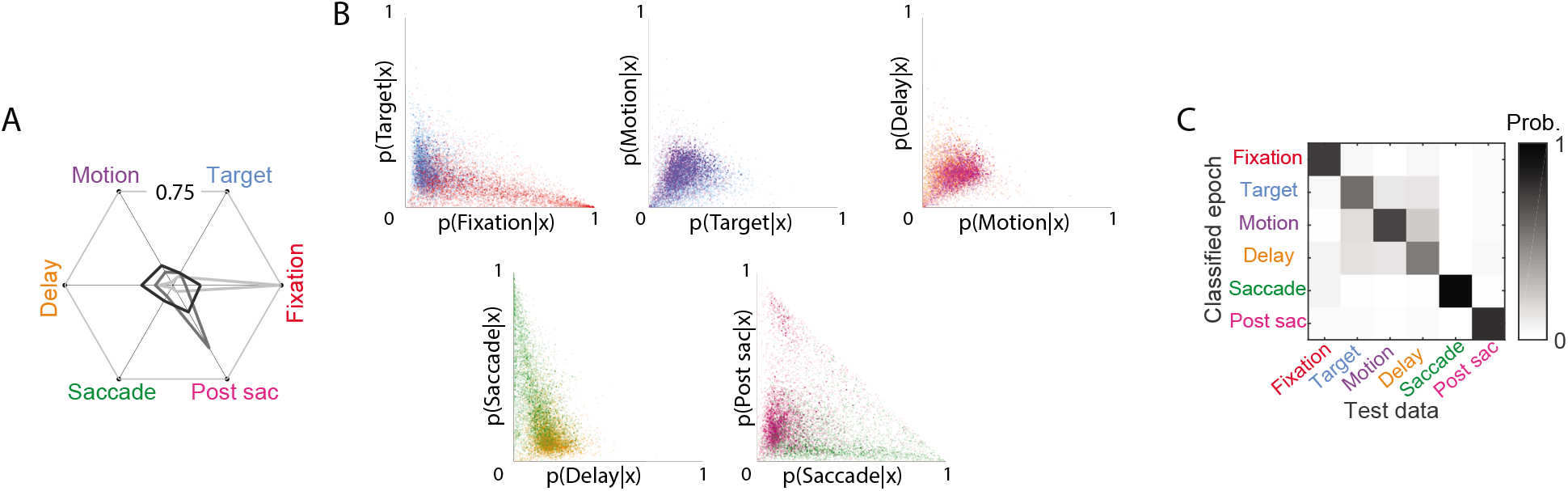
Distinct distributions of population spiking patterns were observed in different task epochs. **A.** The probability of occurrence of 3 example population spike patterns in different task epochs. Each grey polygon connects the probability of a particular population spike pattern to appear in each of the six epochs. Each of these patterns is dominantly present in one of the epochs, but are still likely to appear in the other epochs, albeit with lower probability. **B.** p(epoch|pattern) for all spike patterns occurring in successive epochs of the task. A group of 50 units from a single session is used for the figure. Individual dots represent different spike patterns and their color indicates the epoch in which they appeared. **C.** Epoch classification confusion matrix shows the average likelihood that the observed sequences of spike patterns in an epoch are most likely to come from that or other epochs. Results were averaged over 50 groups of 50 units.

### Distinct vocabularies of population spiking patterns are used in different task epochs

While the same population activity pattern might appear in multiple epochs, their relative occurrence, 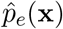, could be vastly different (see examples in Fig. 2A). To investigate how distinguishable the distributions of activity patterns in different epochs are, we compared the likelihood of each epoch given activity patterns, *p*(*e*|*x*), which we estimated using Bayes rule, 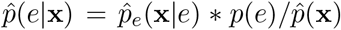. Fig. 2B shows the distinguishability of successive epochs, reflecting that the population activity patterns during the target, motion, and delay epochs were largely similar to one another, whereas the other epochs were more specialized. We quantify the differences between epochs over multiple groups of 50 units in one session using a likelihood-based classifier of activity segments from individual trials. This classifier identified the correct epoch with high accuracy, shown by the decoding accuracy for held out test data in Fig. 2C.

### Temporal dependence and transition rules between population spiking patterns in different epochs

Having mapped the space of population vocabulary, we set out to determine the nature of the trajectories that a neural population traverses, i.e., temporal sequences of activity patterns, and the rules that govern these dynamics. The dynamics of population spiking patterns in different epochs can be described in terms of trajectories in the space of patterns. We show here such trajectories in terms of their likelihood to appear in different epochs (Fig. 3A). To study these trajectories, we extended the pairwise maximum entropy models by adding temporal correlations between units [41, 42]. To model a sequence of states, x(*t – m*), x(*t – m* + 1),…, x(*t*), we extended the maximum entropy framework to find the minimal models that retained the correct firing rates 〈*x_i_*〉 and pairwise correlations at the same time bin 〈*x_i_*(*t*)*x_j_*(*t*)〉, as in eq. (1), but also added the correlations between units at different time bins, 〈*x_i_*(*t*)*x_j_*(*t′*)), for *t* ≠ *t′*. These models give the probability to observe a sequence of *m* + 1 states:

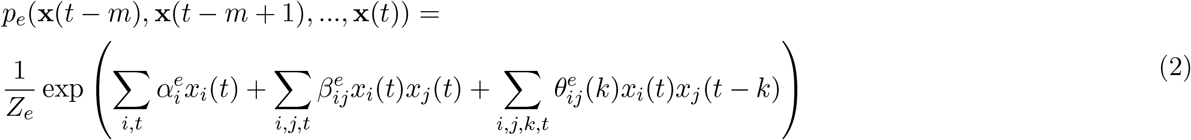

where 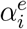, 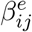, and 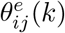 were calculated by fitting the observed population spike patterns over different time gaps, in epoch *e*, and *k* = 1, 2… denotes the duration of gap between the time bins (see a diagram of the interactions between units in the model in Fig. 3B). We compared the performance of different variants of these models, using all possible combinations of parameters: single unit firing rates, pairwise correlations (in the same time bin), temporal autocorrelations of individual units, and pairwise temporal correlations between units. As we discuss below, we found that models that relied on *α_i_, β_ij_*, and *θ_ii_*(*k*), (Fig. 3C) captured most of the spatial and temporal structure needed for highly accurate predictions in all epochs. Using *θ_ij_* (*k*) gave marginally better models in some cases, but suffered from over-fitting issues in others.

**Figure 3:**
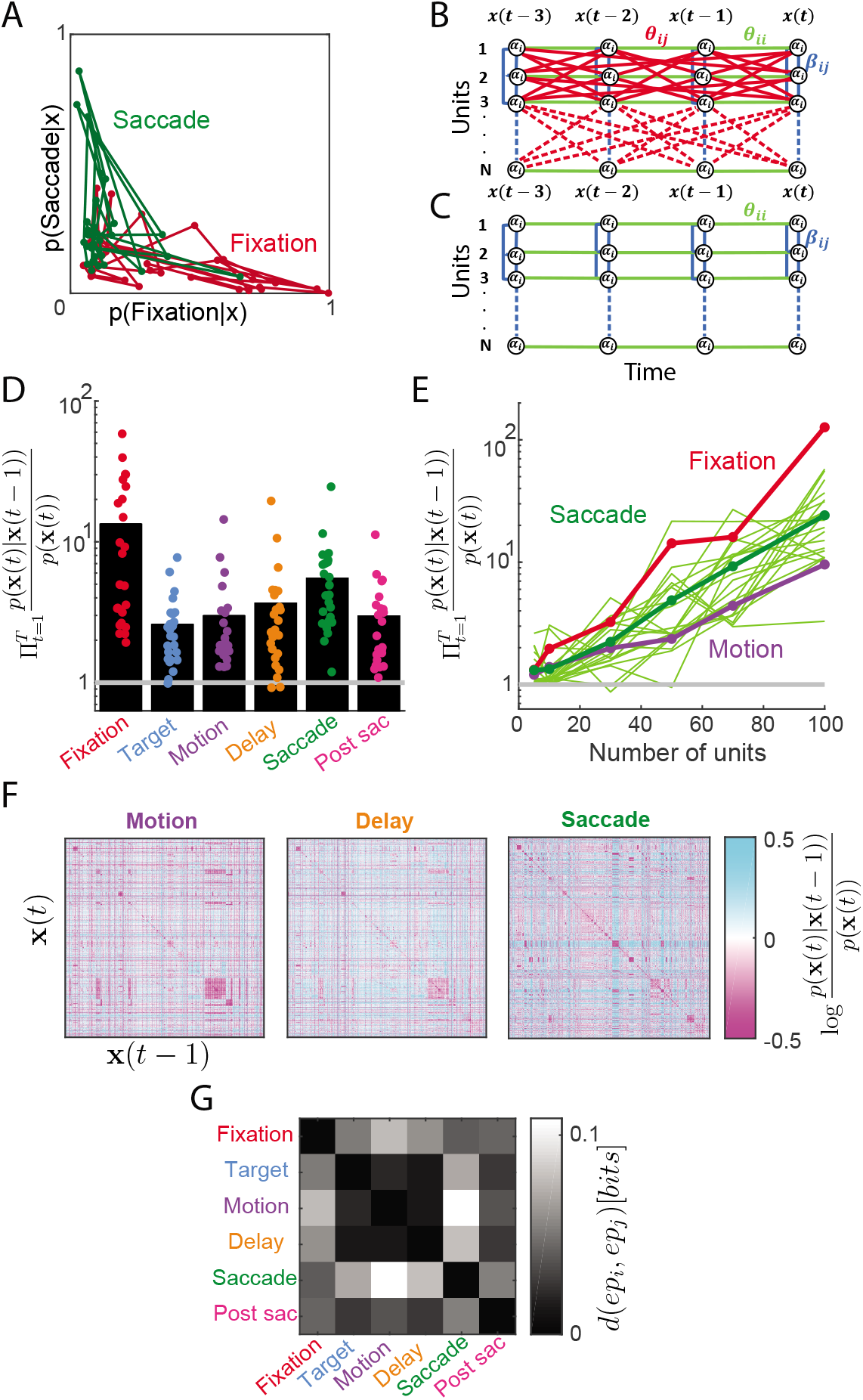
Spatio-temporal models captured the dynamic aspect of the neural code. **A.** For two sequences of population activity patterns from different epochs, we plot the likelihood of different task epochs, p(epoch|pattern). Patterns from the same epoch are connected by a line of the same color based on their sequence. This example reflects the temporal distinguishability of these epochs. **B.** Illustration of the full spatio-temporal maximum entropy model - black circles represent units, with each column representing a different time step. Edges represent interactions, colored by class: spatial interactions, *β_ij_*, are shown in blue, temporal ones between the same cell at different times, *θ_ii_*, in green, and spatio-temporal ones, *θ_ij_*, in red. **C.** Same as in B, for the reduced version of the model, which did not include *θ_ij_*’s. **D.** The ratio between the likelihood of spike pattern sequences in 300 ms epochs under the one time step conditional model *p*(x(*t*)|x(*t* – 1)) and *p*(x(*t*)) for all task epochs. Dots mark individual randomly selected groups of 50 units, and the black bar denoted the average over 20 groups. **E.** Likelihood ratio of temporally conditional models and non-conditional ones as in D, shown as a function of the number of units used for three representative epochs (Fixation, Motion, and Saccade). Bold lines show averages over groups and trials whereas thin lines show individual trials from one session for one of the epochs (Saccade). **F.** Transition probabilities between states are shown for three epochs, for a randomly selected group of 2000 patterns that appeared in the data. Note that in all cases the matrices show the transition probabilities between the same 2000 states in the same order. **G.** Distinguishability of epochs based on the similarity of their state transition matrices (from F) between all pairs of epochs. See main text for details.

From the models of sequences of spiking patterns in eq. 2, we inferred conditional temporal models of the population. These models describe the temporal progression of the population code, namely the probability to observe a population activity pattern following a sequence of patterns, *p_e_*(x(*t*)|x(*t*–1),…, x(*t–m*)). We used these models to quantify the effect of recent history in shaping the neural population trajectories. Our quantification was based on the likelihood ratio of these models and *p_e_*(x(*t*)), namely 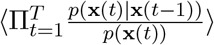. The temporal dependencies were apparent already for a single time bin (*m* = 1) compared to models that ignored temporal correlations: The likelihood ratio of these two models for held out test data significantly exceeded 1 in all task epochs, indicating the effect of recent history on the population activity. The exact magnitude of the effect of history varied across epochs (Fig. 3D). Notably, similar results were obtained for all experimental sessions and monkeys (Fig. S4A). The temporal dependence between states scaled with the size of the population considered (Fig. 3E). We verified that these results were not a consequence of increased model complexity: We generated a similar number of synthetic sequences of population activity patterns from the temporally independent model that was fitted to the data, *p_e_*(x(*t*)), and found that fitting a conditional temporal model to these data *p_e_*(x(*t*)|x(*t*–1)) correctly recovered the lack of temporal correlation in them (Fig. 4B).

**Figure 4:**
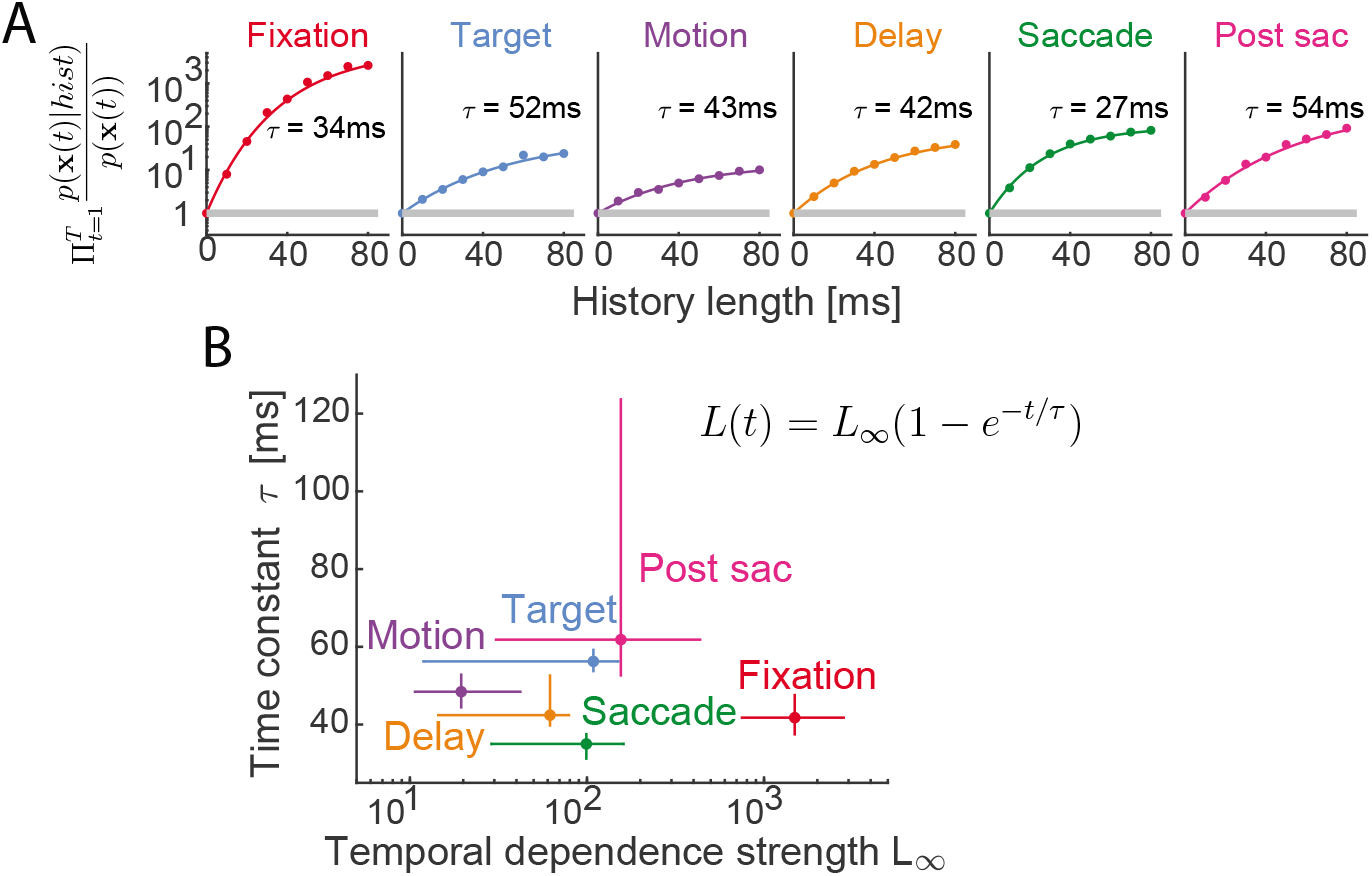
Time scales and strength of history dependence in shaping population spiking pattern sequences. **A.** Likelihood ratio of observed population response trajectories from one session for all rightward choices (sequences of population spiking patterns) of models with different sequence lengths and those of the memoryless models are plotted vs. the length of sequence. Data points represent averaged results over 20 randomly chosen groups of 50 units from one session; error bars representing STD over groups are smaller than marker size. Curves show the fit of an exponential model (see text for details). **B.** Median of the asymptotic values of the exponential fit of the temporal correlations, *L*_∞_, and the corresponding time constants *τ*, are shown for all epochs. Parameters were estimated for 5 sessions separately for leftward and rightward choices in each session; error bars represent the 25 and 75 percentiles.

The conditional models *p_e_*(x(*t*)|x(*t* – 1)) describe the probabilistic “transition rules” between states in each of the epochs. Fig. 3F shows parts of the transition matrices as log *p*(x(*t*)|x(*t* – 1))*/p*(x(*t*)) for three epochs, using the same randomly selected subset of 2000 activity patterns that appeared in held out test data (showing the full transition matrices for all states of a group of 50 units requires 2^50^ rows and columns, which cannot be accommodated in a single figure). These matrices suggest relative similarity in transitions between some epochs (e.g., motion and delay matrices), and clear distinctions between others (e.g. motion vs saccade). We quantified the similarity of the transition matrices between all pairs of epochs *i* and *j* (Fig. 3G), using Jensen-Shannon divergence 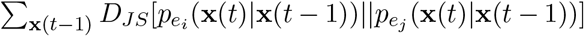. The figure demonstrates that epochs differed not only by the distribution of activity patterns, but also in how patterns transitioned over time. Next, we characterized these transitions for sequences of population spike patterns.

### History-dependence and characteristic time scales of population spiking dynamics in different epochs

The temporal dependence between activity patterns of the population extended beyond the single time step we focused on in the previous section. The likelihood ratio between history-dependent models of the population activity, *p_e_*(x(*t*)|x(*t* – 1),…, x(*t – m*)), and ones that did not use history dependence kept growing with the length of history of states we considered, *m*, for all epochs, as shown in Fig. 4A, for one of the experimental sessions. The improvement with history length did not fully saturate for windows up to 80 ms (which was the maximum length that we could accurately model for groups of 50 units). However, the dependence of the likelihood ratios on *m* were accurately fit with an exponential function of the form *L*(*t*) = *L*_∞_(1 – *e*^-*t*/*τ*^), where the fitted parameters were the time constant for the history dependence, *τ*, and the asymptotic value of the likelihood ratio *L*_∞_.

The differences between epochs were apparent in both the asymptotic values of the correlation effect and their corresponding time constants (typically ranging from 20 to 60 ms), as shown for multiple sessions in Fig. 4B. These values reflect the effect of the temporal correlations in limiting the variety of the neural population trajectories that occur in an epoch. The effect of history in shaping the neural trajectory is strongest during fixation. As might be expected, history has the lowest relative impact on population spiking patterns during the motion-viewing epoch, when randomly varying sensory inputs shape the monkey’s upcoming choice. However, we emphasize that even there, the spiking history limits the set of likely trajectories by many orders of magnitude in likelihood units.

These models reveal the different processing dynamics in the epochs, on scales much shorter than the common temporal windows used for firing rate models. The shortest time scale was found in the saccade epochs (~25 ms), where stereotyped motor plans are executed, whereas the longest time scales were found in target epochs and after the saccade, when the monkey awaited the feedback. The quality of the fits implies that population spiking patterns became conditionally independent of past history after 75-150 ms (~ 3*τ*, which would constitute a temporal Markov blanket [27, 43]).

Comparing the performance of different variants of the maximum entropy models for the epochs reveals the nature of the processing or computation that is performed. We found that the *β_ij_* were critical for the models’ performance (Fig. 5A) such that models without them were orders of magnitude inferior in describing the dynamics of population activity. Adding *θ_ii_*(*k*) captured most of the temporal structure needed for highly accurate models in all epochs (Fig. 5A). Notably, models that also included temporal relations across units *θ*_*i,j*≠*i*_ were better in some sessions but did not generalize on held-out test data for other sessions. Thus, the correlations between units at the same time bin set the baseline structure for the population activity, which is then modulated by the single cell temporal auto-correlations.

**Figure 5:**
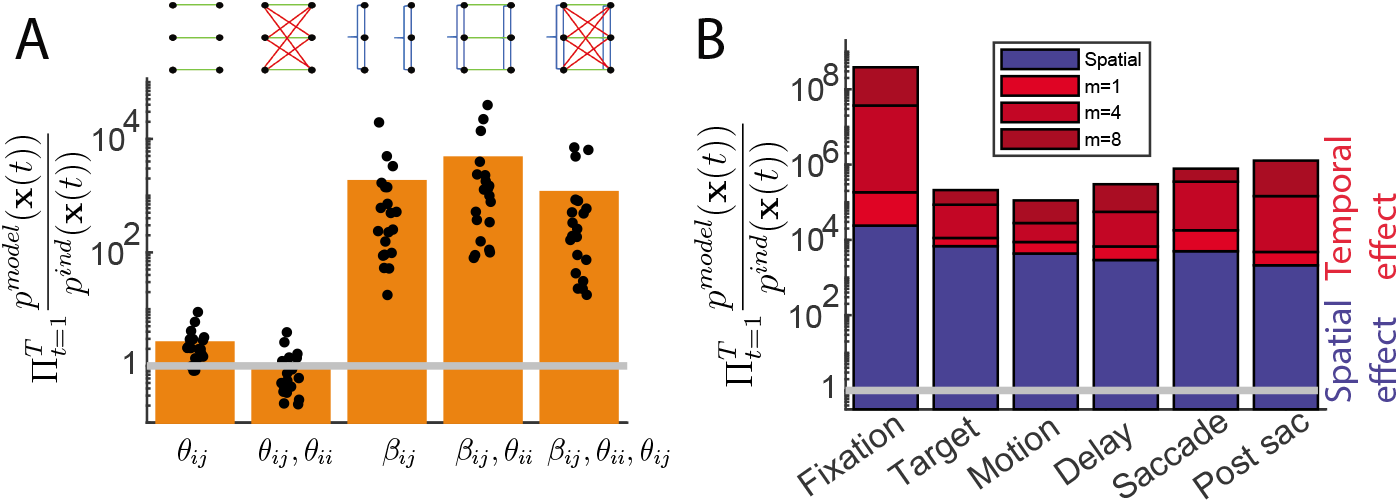
Decomposition of the effect of spatial and long temporal correlations on population activity. **A.** Comparison of spatio-temporal models with different sets of parameters where the reference is the independent model that relies only on firing rates (i.e. models with only *α_i_*’s). Results are shown for one group of 50 cells during the delay epoch of one of the sessions, using a one step history, as a representative example. **B.** Average likelihood ratios of spatial and spatio-temporal models versus the corresponding independent model, for the different epochs. Ratios of spatial models, only with *β_ij_*, are shown in blue, and models also with *θ_ii_* from one step to 8 steps, are shown in shades of red for different history dependence. Data was averaged over 20 randomly selected groups of 50 units.

Fig. 5B summarizes the relative effect of spatial and temporal correlations on the population code in the different epochs. Compared to independent models, the spatial correlations make the observed patterns 10^4^ to 10^5^ times more likely, and the temporal structure increase that by additional 3 to 4 orders of magnitude. Overall, the observed sequence of activity patterns are 10^4^ to 10^7^ more likely than expected from firing rates across different epochs. Similar results were obtained for all experimental sessions and monkeys (Fig. S4). As in the previous sections, using 20 ms time bins did not change the strength and time scales of the spatial or temporal correlations (Fig. S7).

### Temporal sequences of population spiking patterns leading to a decision

Our modeling approach enables us to track when population spiking patterns begin to reflect upcoming choices and how the codes associated with different choices evolve and diverge over time. We fit separate models for each of the epochs - conditioned on the monkey’s choices. Using these models, we calculated the likelihood that an observed spiking pattern would occur for rightward or leftward choices. Assessing the likelihood for a sequence of patterns over a 300 ms window in each epoch, we generated predictions for the most likely choice and compared it with the monkey’s actual choices. The ratio between the likelihood values of models trained on left and right choices in each of the epochs, given by

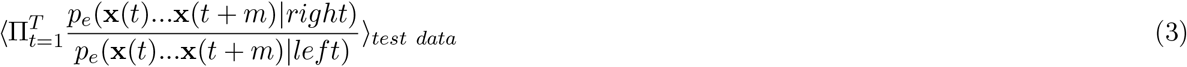

measures the difference between population trajectories for the different choices. Figure 6A shows the likelihood ratios of individual trajectories reflecting how the vocabularies used for right vs left trials become distinguishable during different epochs. Note that we have further divided the Motion epoch into overlapping sub-epochs to better reflect the temporal dynamics of the likelihood ratios. The predicted choices based on these likelihood ratios were highly accurate for the saccade and post-saccade epochs (as expected), and showed a gradual rise during motion viewing and delay epochs (Figure 6B), compatible with formation of decisions based on sensory input in those periods [12, 44–46].

**Figure 6:**
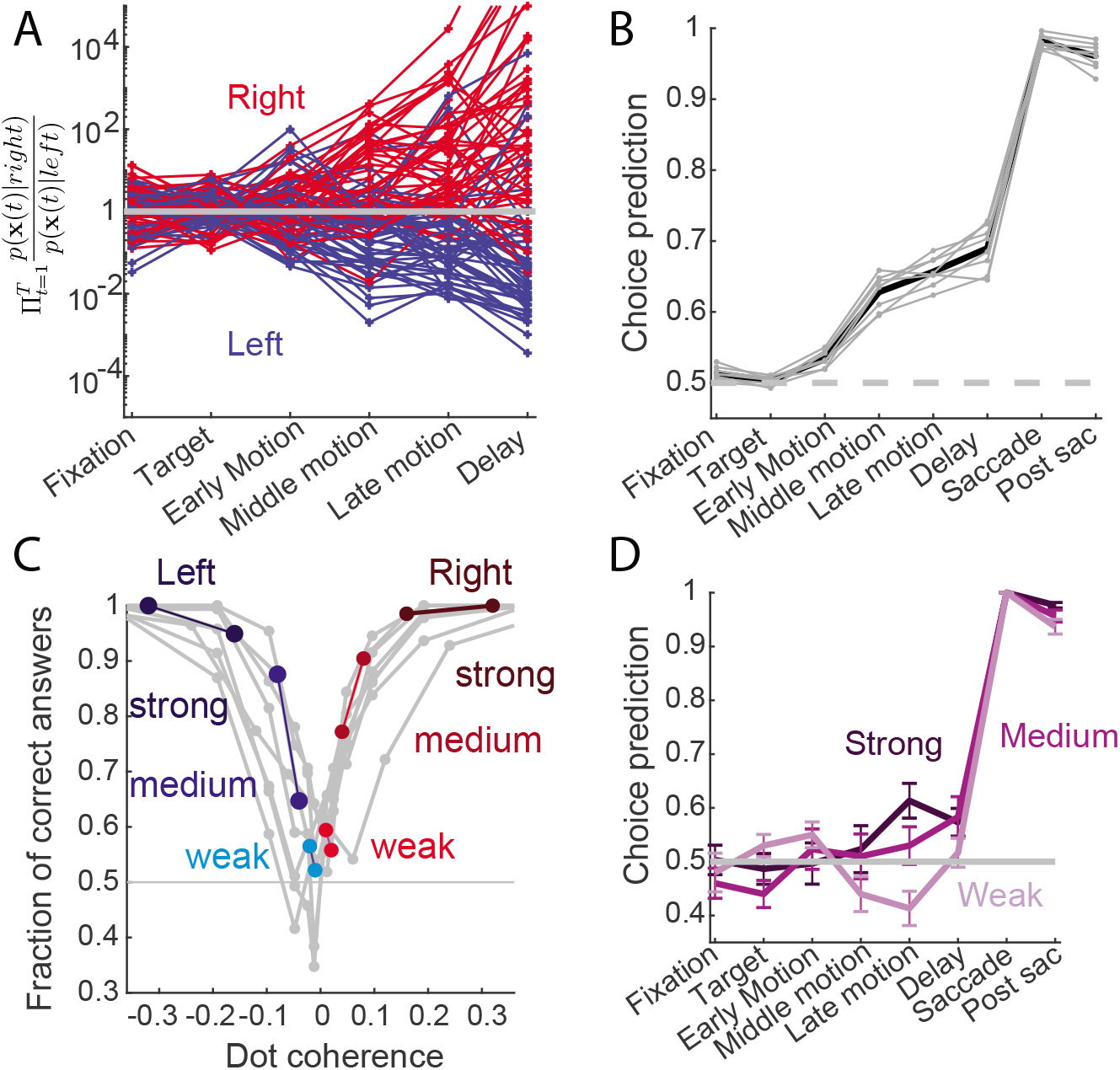
Evolution of neural trajectories towards a decision. **A.** The likelihood of individual trials was computed under a model trained on trials that ended with rightward choices and another model trained on trials with leftward choices. The likelihood ratio of individual test trials is shown for different epochs along the trial. Each line denotes a trial and is colored based on the monkey’s choice. Here, we use three different windows during the motion epoch (early: 100-400 ms after motion onset, middle: 300-600 (this is the same as motion epoch above), and late: 500-800). **B.** Accuracy of decoding the monkey’s decision at different epochs during the trials. Black line shows the average over sessions and gray lines show individual sessions. **C.** Accuracy of monkeys in different experimental sessions is shown as a function of motion coherence. **D.** Accuracy of decoding the monkey’s decision at different epochs along the trials is shown for trials grouped by strength of motion coherence: Weak denotes trials with coherence <5%, Medium, 5-10%, and Strong, >15%.

Because the monkey’s accuracy was dependent on the coherence of the moving dots (Fig. 6C), we investigated how task difficulty influenced the dynamics of population spiking patterns. We split trials into strong, medium, and weak motion strengths, for correct responses to each motion direction, and trained separate models for trials in each group. We found that the difference between left and right choices (Fig. 6D) increased with coherence level (compare e.g. [13]), suggesting that the population spike patterns represent integration of sensory evidence, as previously demonstrated in this brain region [13, 16, 17, 44]. These differences were apparent only motion and delay, and not during saccade, as expected.

### Decoding decisions using different models of population spiking

Complex encoding mechanisms and correlations can result in codes that are easy to decode, and simple encoding might result in codes that are hard to decode. This idea has been common in theoretical models, such as liquid state machine [47, 48] and decoding algorithm as in [49], Hopfield networks [50], adaptive codes [6], predictive coding [51–53], as well as other biological systems such as genetic networks. In particular, evaluating encoding by a certain class of decoders can give useful lower bounds on the amount and nature of encoded information, but may not give the correct interpretation of the encoded information.

Following these ideas, we asked how well we can decode the monkey’s choices at different epochs, using the independent models and the pairwise models based on spatial and temporal correlations between units. Despite the strong impact of spatial and temporal correlations on the neural population responses and dynamics, we found that the independent models were largely on par with the best pairwise models for decoding the choices (Fig. 7).

**Figure 7:**
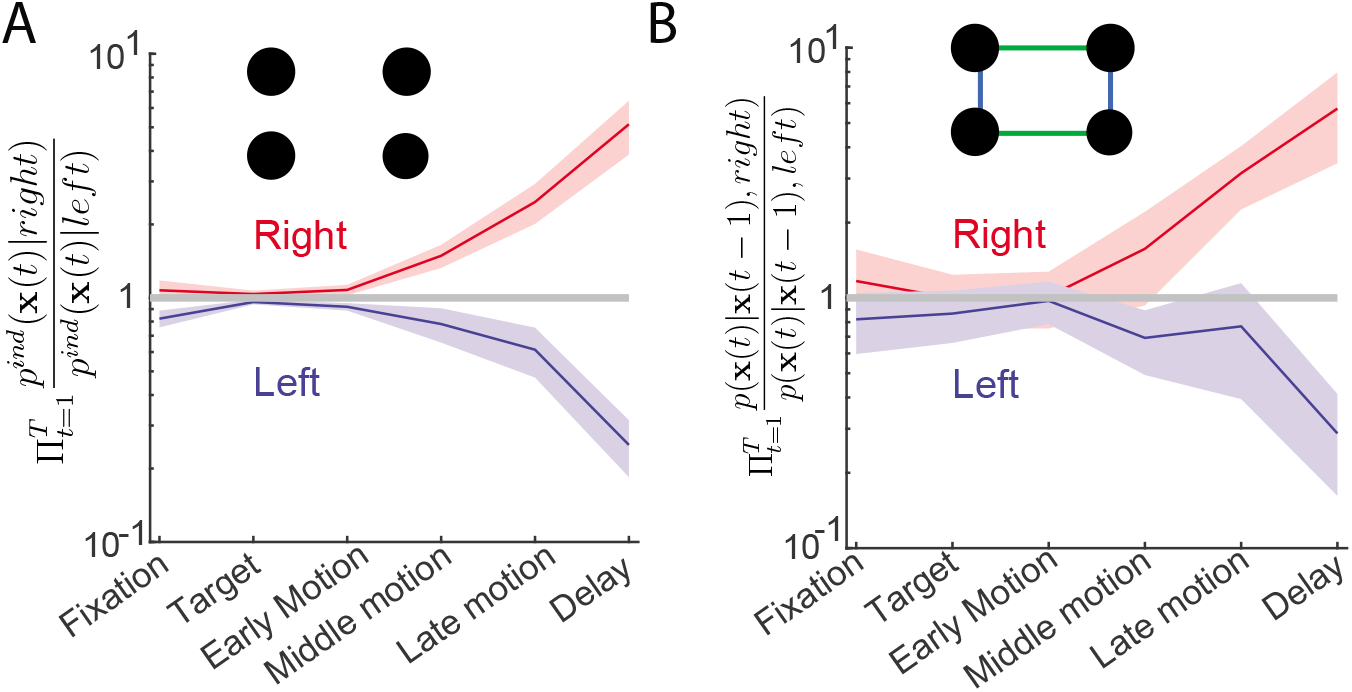
Decoding behavior using correlations and without them gives similar results, despite the strong network correlations. **A.** The likelihood ratio of independent models for right and left decisions was compared for each epoch, on trials that ended with rightward and leftward decisions. Results shown are averages over individual trials in different sessions. **B.** Same as (A), using the spatio-temporal models.

The independent model has a much lower number of parameters (*o*(*n*) instead of *o*(*n*^2^) for the pairwise model) and is, therefore, easier and faster to train with limited samples. The observation that the decoding accuracy of the independent and best pairwise models are largely similar partially stems from finite experimental data. However, it also demonstrates that while correlations are essential for encoding dynamics of the prefrontal cortex, they are not the only information bearing elements in the neural code. Rather, the prefrontal code is accessible to simple linear decoders that do not need to know details of local circuitry and correlated activity in the population to be accurate.

## Discussion

We have presented a detailed analysis of the spatio-temporal structure of the code of large populations of neurons in the prefrontal cortex of monkeys performing a perceptual decision-making task. This is the first time that the relative contributions of the spatial and temporal structure of the spiking patterns of large populations in the primate prefrontal cortex and their dynamics have been mapped at a fine resolution in individual trials. By using the maximum entropy framework to build the mathematically minimal statistical models of the data that rely on different correlation structures, we were able to quantify the effect of firing rates, spatial correlations, and temporal correlations on the neural population vocabulary in different epochs of individual trials and their dynamics.

We found that the correlations among units have a very strong impact on the vocabulary of large populations and their dynamics. The observed vocabulary is many orders of magnitude more likely to appear under models that incorporate correlations between units. Spatial correlations (over a short time window of 10 ms) have the strongest impact, and together with the temporal correlations over tens of milliseconds, cause the spike pattern sequences observed in the data to be 10^4^ to 10^7^ times more likely than predicted by models that ignore the correlations and rely just on the firing rates of individual units. Thus, the population code of the prefrontal cortex is shaped by spatio-temporal correlations. These models reveal not only the strength of correlations in shaping the code for different task epochs, but also the characteristic time constants for the dynamics of sequences of states in different epochs of the task. We found these time constants to be generally shorter than the typical temporal windows used in previous studies.

A similarly strong impact of correlations for encoding of information by large populations of neurons has been observed in different neural systems, from the retina, to visual cortex, hippocampus, and the prefrontal cortex. While these systems differ in the nature of the dominant correlations and their temporal structures, the unifying theme is one of population codes that are shaped by the correlation structures. The implications are a codebook and sequences of states that inhabit a much narrower subspace of the high dimensional population activity state space. Binary spike patterns occupy corners of a hypercube in the native state space of population activity. Correlations in the neural code limit the visited corners and dynamics of transitions from one corner to others. One can think of “tubes” embedded in a higher dimensional space that define the set of possible transition vectors in the state space. In the population code of PFC, the “width” of these tubes is much narrower and dynamics within them much more directed than expected from a population of independent units. Past theoretical analyses have suggested that such designs would maximize information transmission in noisy environments, and help match the neural code to the statistical structure of relevant stimuli [54].

Given the prevalent and dominant nature of correlations in the code of large populations, it might seem surprising that decoding of information from neural populations that ignore them has been so successful. We have found that ignoring the correlations in decoding, results in performance which is roughly on par with decoders that take them into account. While this may be an idiosyncratic feature of our task (see e.g. [55]), we suggest that this may also be a design principle of the code: While encoding depends on the dynamics of the inputs and the synaptic organization of the circuit, these complicated correlated codes may have evolved to be read easily. In particular, a decoder that ignores correlations that have a huge impact on encoding would still give very accurate decoding. The idea of complicated encoding systems yielding easily decodable codes has been previously suggested [56, 57]. Although simple decoders have been successfully used to read out neural population responses in diverse cortical circuits [13, 15, 58–62], the complexity of the neural encoding and decoding has been rarely quantified in the same dataset. We provide such a direct comparison in large populations of simultaneously recorded cortical neurons for the first time, demonstrating strong influence of history-dependent spatial and temporal correlations on encoding in conjunction with a simple decoding scheme.

Our results match previous studies in the retina, where it has been shown that much of the semantic organization of population codes can be learned by models that ignore correlations, despite the orders of magnitude failures of such independent models in capturing the encoding dictionary. This organization of the code facilitates using linear decoding as an efficient means for learning and reading out behaviorally relevant information [35, 36]. The match between retinal and prefrontal populations, as well as similar results in sensory cortex [32], suggest common functional and design principles that govern encoding and decoding across biological neural circuits.

Recent analysis of history dependence in the posterior parietal cortex [21], has shown correlations in population activity in mice performing a spatially and temporally extended decision making task, which lasted over seconds. Aside from differences in the species and the nature of the task, our modeling of spiking patterns focused on much shorter time windows and allowed us to characterize the underlying temporal structure of dependence between population states. In particular, the time scales we identified reflect the windows beyond which population activity becomes independent of its past, even if correlated. In other words, one can get arbitrarily long temporal correlations where the intrinsic window of dependency is much shorter. Linking the time scales in [21] and the population spiking time scales we find here would hopefully enable the dissection of computation and “working memory” in the PFC.

Our analysis of population dynamics through modeling of spiking patterns using fine temporal windows, brings together the power of decomposition of spatio-temporal structure of population activity and detailed modeling of spiking patterns of large neural populations. This combination extends methods like tensor decomposition [63, 64], as we provide an interpretable generative model and do not depend on metric based approaches, and machine learning based approaches for modeling spiking patterns as autoencoders (LFADS). More broadly, the current framework is easily applicable to other neural systems, and can be used to study the nature of sequences of states [18, 19, 65].

## ACKNOWLEDGMENTS

We thank Roy Harpaz, Tal Tamir, Adam Haber, Yoni Mayzel, Omri Camus, Benny Brazowski, and Gasper Tkacik, for discussions and suggestions. This work was supported by a CRCNS grant (ES and RK), a Simons Collaboration on the Global Brain grant 542997 (RK and ES), a European Research Council Grant 311238 (ES), an Israel Science Foundation Grant 1629/12 (ES), research support from Martin Kushner Schnur and Mr. and Mrs. Lawrence Feis (ES), and Pew Scholarship in Biomedical Sciences (RK).

## Methods

### Experimental setup

Monkeys performed a direction discrimination task with random dots. Each trial began with fixating on a fixation point (0.3^*o*^ diameter) at the center of the monitor. Eye position was measured with a scleral search coil or an infrared tracking camera. Once the monkey fixated, two targets appeared on the screen, indicating the two possible motion directions. Afterwards, a random dots stimulus was shown for *T_stim_* = 800*ms* [13, 37]. Monkeys were trained to discriminate whether the net motion direction was toward one target or the other. The proportion of dots moving coherently in the same direction (coherence or stimulus strength) determined task difficulty, and was randomly chosen in each trial from a fixed set. Stimulus offset was followed by a delay period of variable duration usually ranging from 300 ms to 1500 ms. The delay period ended in a ‘go’ cue, marked by the disappearance of the fixation point, after which the monkey made a saccade to one of 2 targets. If the chosen target matched the direction of motion, the monkey received a liquid reward. A distinct tone was played for errors. Trials with 0% motion coherence were rewarded randomly with a probability of 0.5.

### Experimental Data

Extra-cellular recordings were performed from neural populations of the prefrontal cortex of macaque monkeys. All experimental procedures conformed to the National Institutes of Health Guide for the Care and Use of Laboratory Animals and were approved by the New York University Animal Welfare Committee and Stanford University Animal Care and Use Committee. Recordings from the prefrontal cortex were obtained by implantation of 96-channel Utah arrays in the prearcuate gyrus (area 8Ar) of macaque monkeys (macaca mulatta). During the experiments, monkeys performed a direction discrimination task with random dots [13, 37]. Neural spike waveforms were saved online (sampling rate, 30 kHz) and sorted offline (Plexon Inc., Dallas). Throughout the paper we use the term “units” to refer to both well-isolated single neurons and multi-units. Number of units in different sessions ranged from 169 to 250 units.

### Definition of task epochs

Task epochs were defined using experimental behavior and neural activity (see Fig. S1B), as in earlier studies [13]. We chose 300 ms windows in each epoch consisting of 30 bins of Δ*t* = 10*ms*, or 15 bins of Δ*t* = 20*ms*. In learning the spatio-temporal models, which require history dependence, we used up to 100 ms preceding the analysis window as the history of neural activity for the first time point. Epochs were defined as follows: *Fixation* - starting 100 ms after fixation point onset; *Target* - starting 100 ms after target onset; Motion - starting 300 ms after dots onset; *Delay* - starting 100 ms after dots offset; *Saccade* - starting 150 ms before saccade onset; *Post-saccade fixation* - starting 250 ms after eyes reached the chosen target.

### Modeling population activity patterns using maximum entropy models

As described in [40], the maximum entropy framework gives the most parsimonious or random statistical model, given a set of observable functions of the data. As we discretized neural activity into small temporal bins of size Δ*t*, which were either 10 or 20 ms long, the spiking or silence of neuron i at a particular time bin was given by a binary variable *x_i_* (‘1’ denotes spiking in that bin, and ‘0’ denotes silence) and the binary vector x designates the population activity pattern at that time bin. For a set of observable functions of the form 〈*f_μ_*(x)〉 (where 〈〉 denote the average over the data), the maximum entropy model for that set of observables or constraints is given by finding 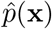 that maximizes

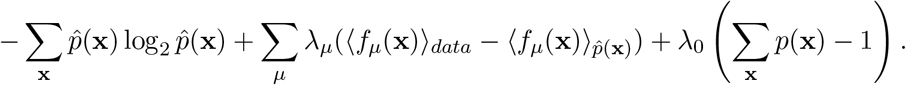

This distribution is given by 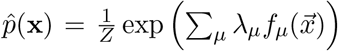, where the Lagrange multipliers, λ_*μ*_ are found numerically, and *Z* is a normalization term or partition function. If the observables are only the firing rates of individual units, 〈*x_i_*〉, then the model is given by 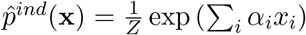, or the independent model. Adding the pairwise correlations between all neurons, means adding constraints for 〈*x_i_x_j_*〉, and the pairwise maximum entropy model is given by 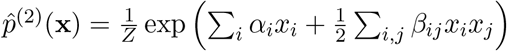 (see also [25, 26]). Adding synchrony constraints gave the k-pairwise models [31]. Spatio-temporal models added observable functions of the form 〈*x_i_*(*t*)*x_j_*(*t* + *τ*))_*t*_, and the form of their models is given in equation 2 (see also [34, 54]). The random projection models rely instead on random nonlinear observable functions as described in [32].

Models were fitted as described in [32], using the maxent toolbox [66], except for the parameters of the independent models that were found using its closed-form solution. The code used to train the models is publicly available [66] as an open-source MATLAB toolbox at https://orimaoz.github.io/maxent_toolbox/. Briefly, we trained the probabilistic models by seeking the parameters λ_*μ*_ that would minimize the Kullback-Leibler divergence between the model 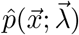 and the empirical distribution 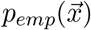, which is equivalent to maximizing 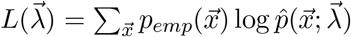, which is a concave function whose gradient is given by

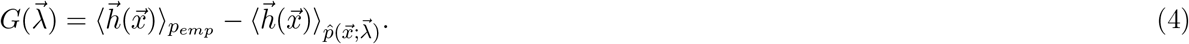

We found the values λ_*μ*_ that maximize the likelihood by iteratively applying the gradient (Eq. 4) with Nesterov’s accelerated gradient descent algorithm. We computed the empirical expectation in Eq. 4 (lefthand term) by summing over the training data, and the expectation over the parameters 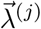 by summing over synthetic data generated from 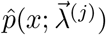 using Metropolis-Hasting sampling.

### Measuring similarity of coding dictionaries

We quantified the difference between coding vocabularies using the Jensen-Shannon divergence *D_JS_*[*p*(x)|*q*(x)] [3], a symmetric measure of the similarity between two probability distributions *p*(x) and *q*(x), which equals 0 if the distributions are identical, and 1 for non-overlapping distributions. It is defined as

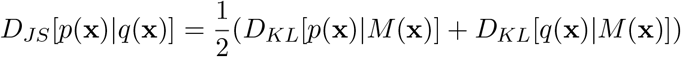

where 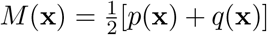, and *D_KL_* is the Kullback-Leibler divergence [3], given by 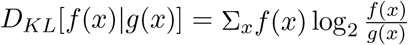, a non-negative measure, which equals 0 if the distributions are identical.

