## Supplementary material for "Strongly correlated spatiotemporal encoding and simple decoding in the prefrontal cortex"

**Recording sites.** The location of the electrodes in the prearcuate gyrus is shown in Fig. S1A, relative to the principal and the arcuate sulci. Fig. S1B displays the mean firing rates and correlations, as well as experimental markers, that guided us in selecting the task epochs. Fig. S1C displays the likelihood ratio between the pairwise and independent models in estimating a sequence of states that appear in an epoch, same as in Fig. 1E, shown for all 15 data sets from both monkeys.

**Comparing the performance of different population models.** Throughout the paper we focused on pairwise models, but we also explored the extended, more potent K-pairwise [1] and random projection models [2]. Fig. S2A shows the likelihood ratio of all 3 population models to the independent model. Based on their similarity, we used the pairwise model, because of its simplicity and interpretability. Fig. S2B shows the performance of all population models on estimating triplet co-spiking $\langle x_i x_j x_k \rangle$ of units accurately, where the independent models make large errors. Fig. S2C shows the level of synchrony in the population, by the probability for $K$ units to spike in the same time bin.

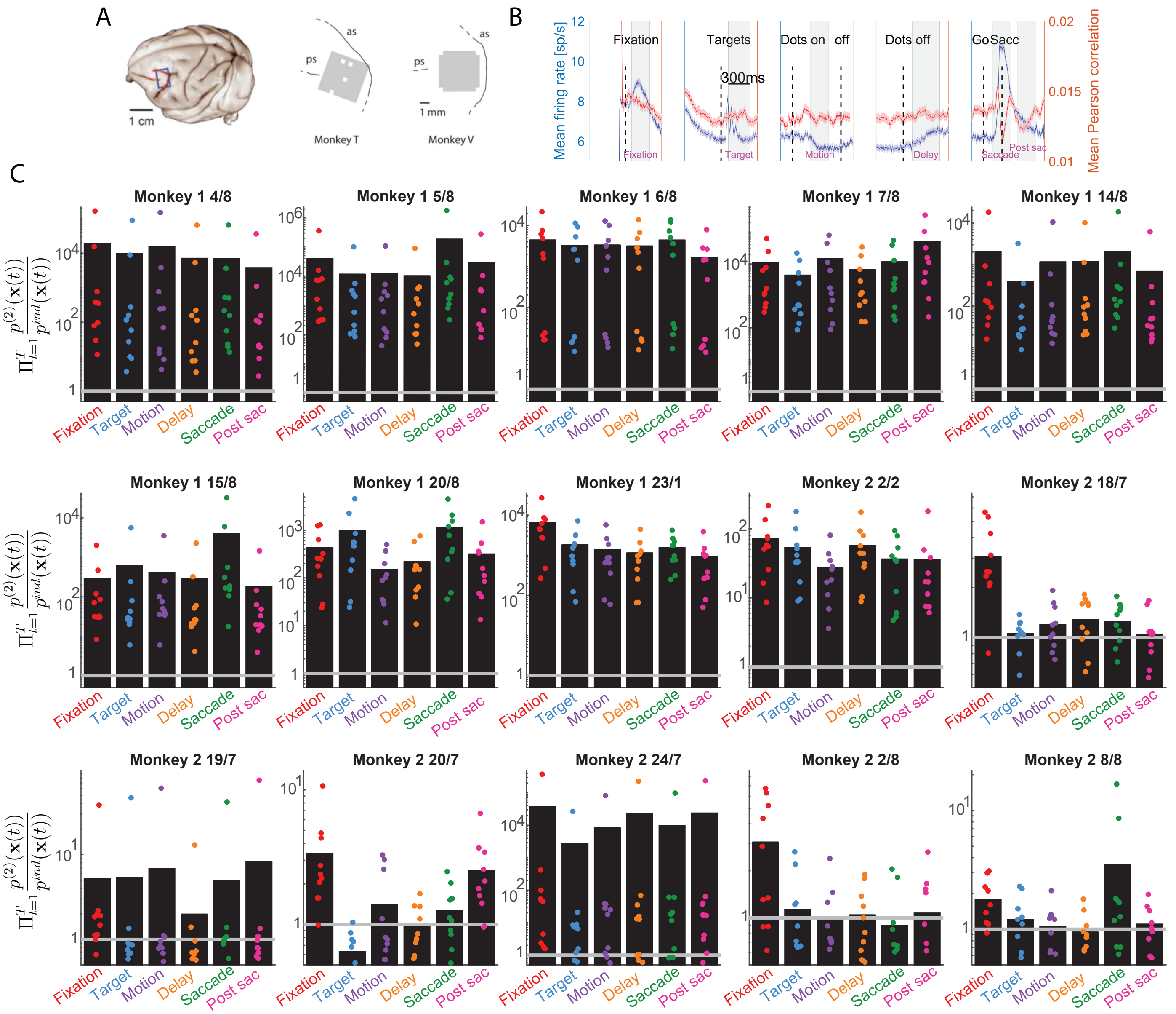


Figure S1: **Correlated activity between of prearcuate gyrus neural populations for multiple data sets. Related to Fig. 1**. **A.** Region of electrode position, in area 8Ar of the prearcuate gyrus (left panel). The two right panels show an accurate position of the electordes for both monkeys, relative to the arcuate sulcus (AS) and the primary sulcus (PS). White squares represent locations of the ground pins. **B.** Firing rates (blue curve), averaged over all units and all trials, vs. time in the trial, as well as Pearson correlation, averaged over all pairs and all trials. Experimental markers are plotted as dashed vertical lines, and epochs used throughout the paper are marked by gray rectangles. **C.** Likelihood ratio of pairwise and the independent models for the sets of states that appear in an epoch (as in figure 1F), shown for all 15 sessions from both monkeys.

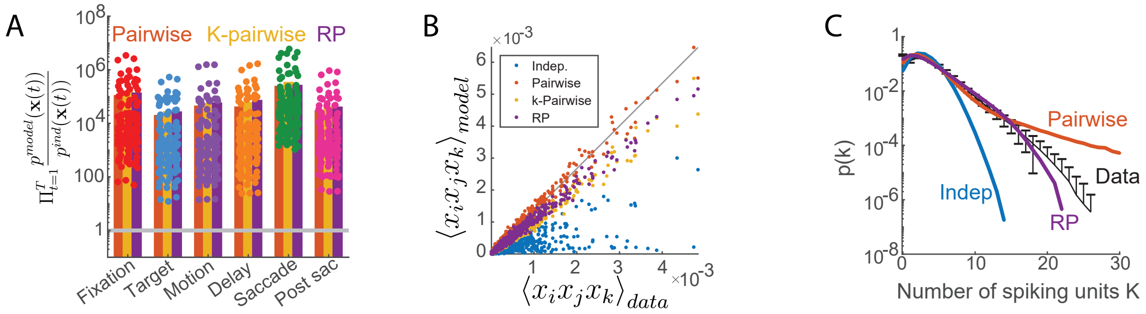


Figure S2: **Comparing different population models. A.** Likelihood ratio of different population models to the independent model. Results are averaged over 50 groups of 50 units, and each dot corresponds to a single group. **B.** Triplet co-spiking probability $\langle x_i x_j x_k \rangle$ as predicted by the models, vs. that in the test data, for groups of 50 units. 1000 randomly chosen triplets from 50 groups are shown, with black line marking equality. **C.** Population synchrony, which is the probability to observe exactly $K$ co-spiking units, as predicted by the models, and as found in the test data, for groups of 50 units.

Fig. 2C in the main text showed the decoding accuracy for groups of 50 units using one of the sessions. In Fig. S3A we show the monotonic increase of the mean decoding accuracy as a functions of the number of units, and obtain $> 0.8$ accuracy for the largest group sizes we studied. We show in Fig. S3B the mean decoding accuracy for all 15 sessions, which is well above chance in all data sets; notably, the decoding accuracy from data sets of monkey T was higher. Fig. S3C shows the confusion matrix for all data sets (see Fig. 2C), and we find in all cases high success rate for the fixation, saccade, and post saccade epochs, and higher confusion rates in the intermediate epochs.

**Performance of spatio-temporal models.** In Fig. S4A we show the likelihood ratio of occurrence of a sequence of states that appeared in an epoch, as predicted by the spatio-temporal vs. the spatial models $\Pi_{t=1}^{T} \frac{p(\mathbf{x}(t)|\mathbf{x}(t-1))}{\mathbf{x}(t)}$, as in Fig. 3D, for all 15 data sets. We notice similar behavior in all data sets, though results for monkey T display a stronger temporal structure. The spatio-temporal models have a large number of parameters, and to control for over-fitting, we analyzed the likelihood ratio between models using synthetic data, for which we know the "ground truth" temporal structure (Fig. S4B). While there is a clear advantage for the spatio-temporal model in the data, for identical analysis of synthetic data that is temporally independent - the models show similar performance. Thus, the advantage of the spatio-temporal models is indeed a result of the structure of the data, and not of our fitting procedure. For temporally-dependent synthetic data, such that the state at time $t$ is generated from the conditional distribution $p(\mathbf{x}(t)|\mathbf{x}(t-1))$ we observe similar results to the real data.

**Extended temporal dependence of the population code.** Fig. 4 showed the improvement in model accuracy with history dependence for one of the sessions. In Fig. S5A we show similar results for several data sets from both monkeys. Training these long-memory models was computationally demanding, and so we show here results for six of the sessions. To control for over-fitting due to the large number of parameters in the model, we again generated synthetic data where we know the full temporal structure. As expected, the corresponding curves in the synthetic data level off at correct memory length with which the data was generated, indicating the results for the real data reflect its true structure, and not due to over-fitting.

Fig. 5B showed the relative contribution of the spatial correlations and the temporal correlations of several orders, in capturing the structure of population activity in the epochs. In Fig. S6 we show similar results for the data sets for which we fitted these long-memory models.

**Models' performance for different temporal bin size.** All results presented in the main text used $\Delta t = 10ms$ time bins. To verify our results do not depend on the choice of temporal resolution, we analyzed the same data using $\Delta t = 20ms$ bins. The effect of the spatial correlations (Fig. S7A) is similar to that shown in Fig. 1E. Similarly, the temporal correlations have a similar qualitative behavior, as observed by

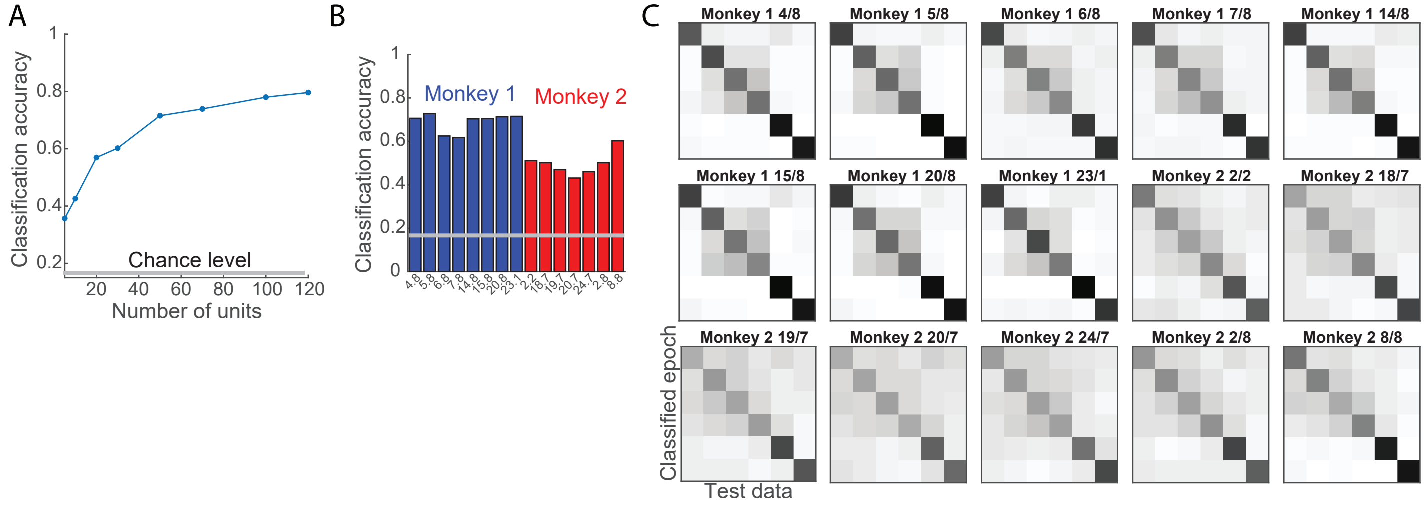


Figure S3: **Epoch classification for multiple sessions and different group sizes A.** Mean decoding accuracy between epochs, plotted vs. the number of units. Results were averaged over 20 groups using spatial models. **B.** Mean decoding accuracy for all 15 sessions, averaged over 20 groups of 50 units. **C.** Confusion matrix for epoch classification (see Fig. 2D) for all 15 sessions.

comparing Fig. S7B to Fig. 3D. Extending this analysis to longer memory models, we found similar increase of model performance with memory, when using 10 and 20 $ms$ time bins (Fig. S7C), indicating the time constants we estimated reflect the true temporal structure of the code.

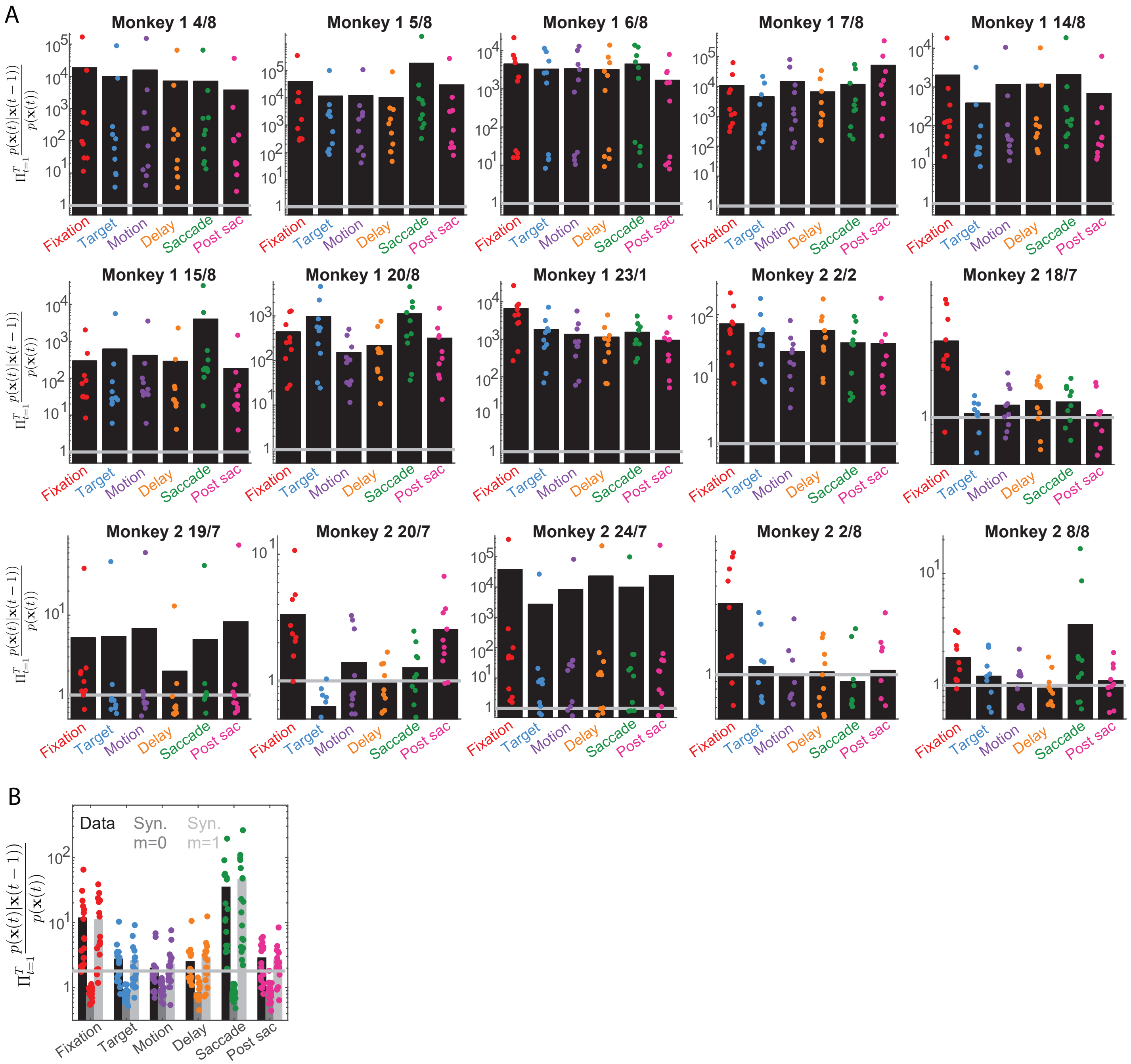


Figure S4: **Comparison of spatio-temporal and spatial models for different sessions. A.** Likelihood ratio between the spatio-temporal and the spatial models for a sequence of states appearing in one epoch (see Fig. 3D) for all 15 data sets. **B.** $\Pi_{t=1}^{T} \frac{p(\mathbf{x}(t)|\mathbf{x}(t-1))}{p(\mathbf{x}(t))}$ computed for the data (as in Fig. 3), for synthetic data without temporal correlations, and for synthetic data where states depend on the past time step. In A,B bars display averaged over groups, and the values of each group are displayed as dots.

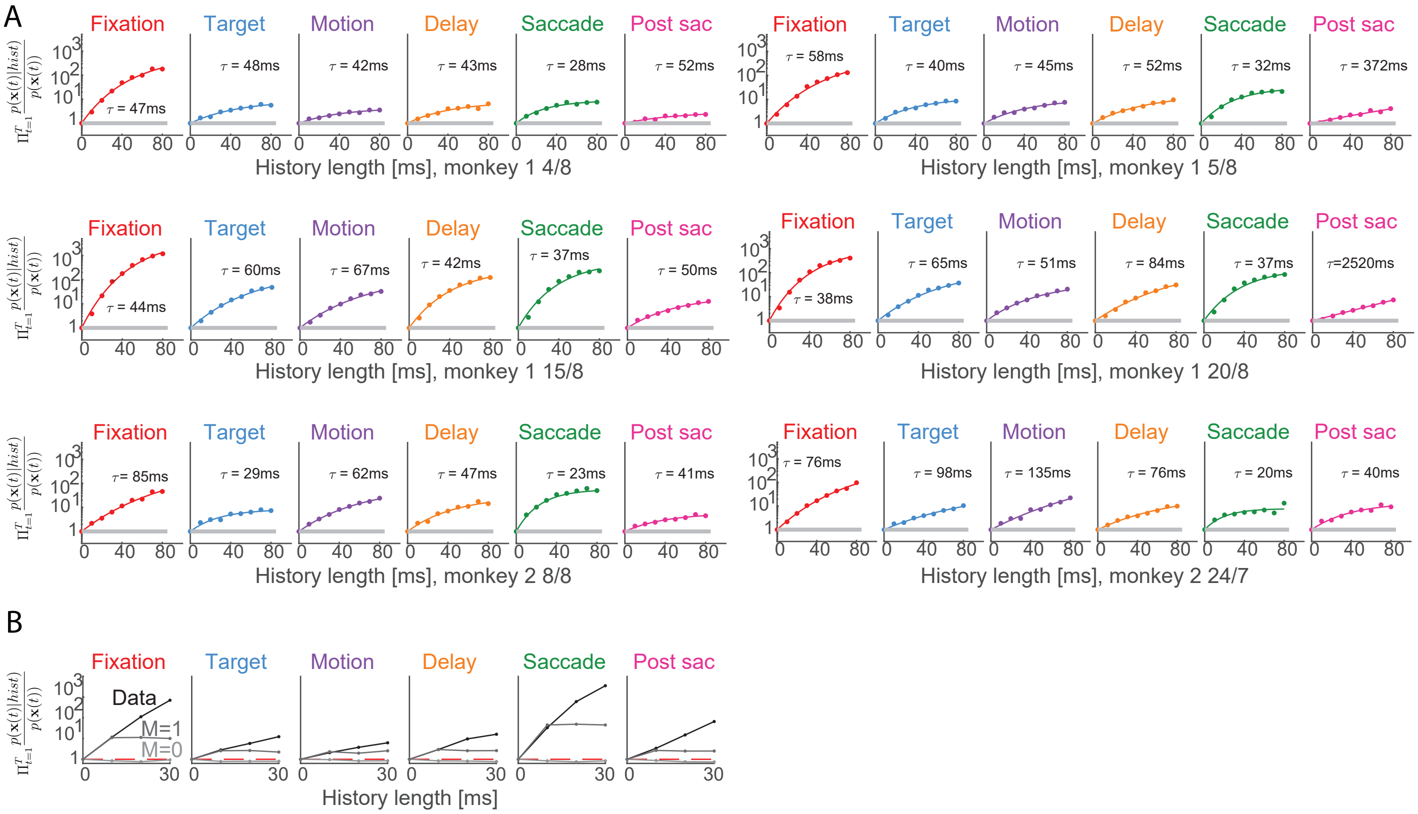


Figure S5: **Spatio-temporal models with long memory for different sessions. Related to Fig. 4 A.** Improvement of models with memory (same as Fig. 4A) for different data sets from the same monkey, and for a data set from the other monkey. **B.** Same as in A, for the data, for synthetic data which is temporally independent, and for synthetic data which has a 1-step memory.

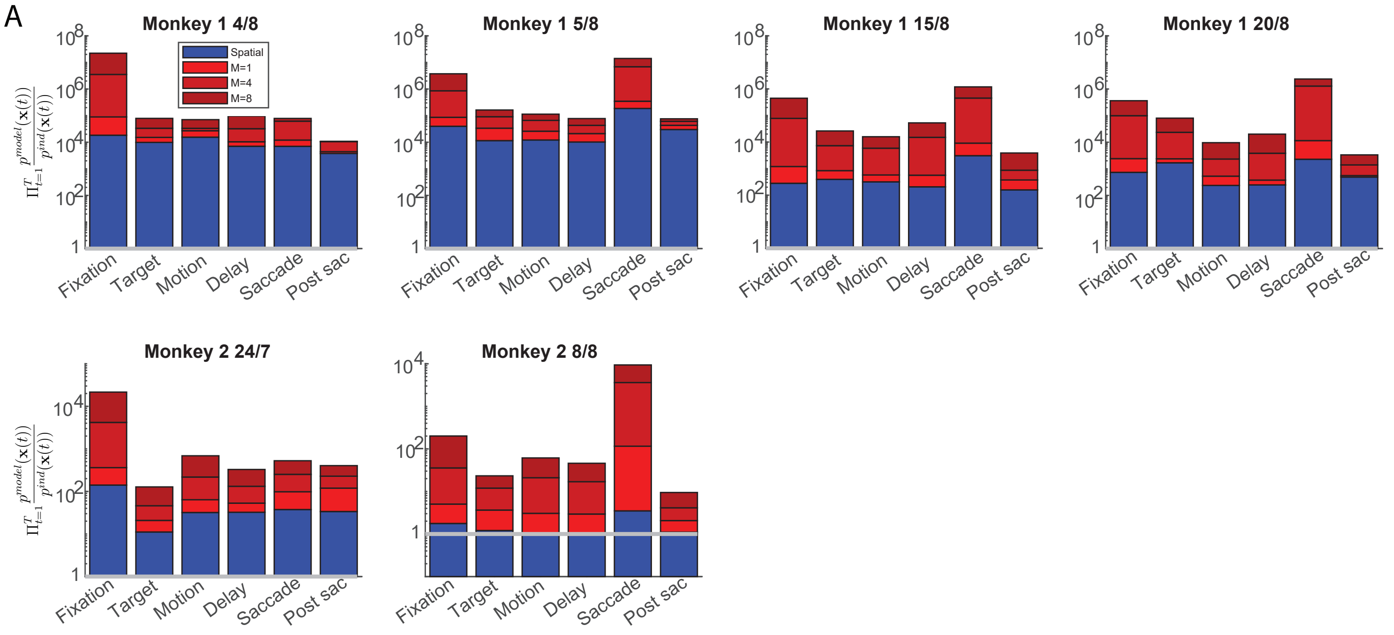


Figure S6: **Spatio-temporal models with long memory for different sessions. Related to Fig. 5 A.** Likelihood ratio for a sequence of states for different models relative to the independent model (see Fig. 5B) for several data sets from both monkeys.

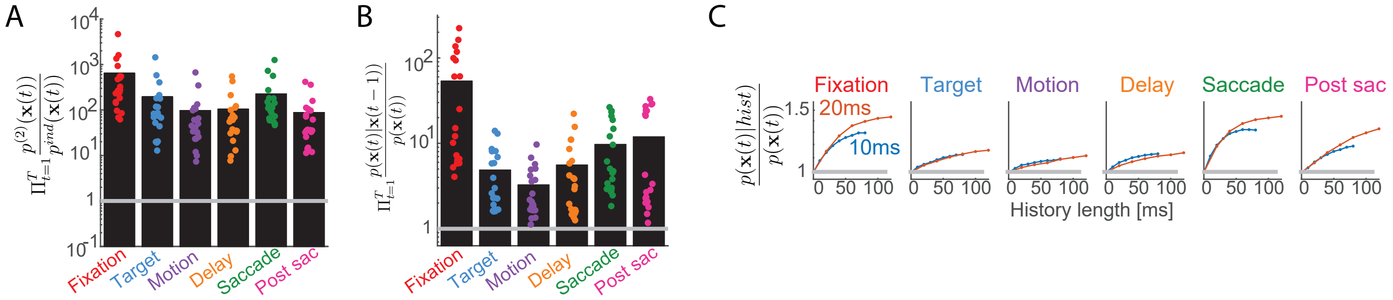


Figure S7: **Model comparison using $\Delta t = 20ms$ time bins.** **A.** Likelihood ratio between the pairwise and independent model (see Fig. 1E) using 20 $ms$ time bins. **B.** Likelihood ratio between a spatio-temporal and a spatial models (see Fig. 3D) using 20 $ms$ time bins. **C.** Likelihood ratio vs. length of memory of the models (see Fig. 4A) when using 20 $ms$ time bins.
